# Intra-species transcriptional profiling reveals key regulators of *Candida albicans* pathogenic traits

**DOI:** 10.1101/2021.03.03.433840

**Authors:** Joshua M. Wang, Andrew L. Woodruff, Matthew J Dunn, Robert J. Fillinger, Richard J. Bennett, Matthew Z. Anderson

## Abstract

The human commensal and opportunistic fungal pathogen *Candida albicans* displays extensive genetic and phenotypic variation across clinical isolates. Here, we performed RNA sequencing on 21 well-characterized isolates to examine how genetic variation contributes to gene expression differences, and to link these differences to phenotypic traits. *C. albicans* adapts primarily through clonal evolution and yet hierarchical clustering of gene expression profiles in this set of isolates did not reproduce their phylogenetic relationship. Strikingly, strain-specific gene expression was prevalent in some strain backgrounds. Association of gene expression with phenotypic data by differential analysis, linear correlation, and assembly of gene networks connected both previously characterized and novel genes with 23 *C. albicans* traits. Construction of *de novo* gene modules produced a gene atlas incorporating 67% of *C. albicans* genes and revealed correlations between expression modules and important phenotypes such as systemic virulence. Furthermore, targeted investigation of two modules that have novel roles in growth and filamentation supported our bioinformatic predictions. Together, these studies reveal widespread transcriptional variation across *C. albicans* isolates and identify genetic and epigenetic links to phenotypic variation based on co-expression network analysis.

**Importance:** Infectious fungal species are often treated uniformly despite clear evidence of genotypic and phenotypic heterogeneity being widespread across strains. Identifying the genetic basis for this phenotypic diversity is extremely challenging because of the tens or hundreds of thousands of variants that may distinguish two strains. Here we use transcriptional profiling to determine differences in gene expression that can be linked to phenotypic variation among a set of 21 *Candida albicans* isolates. Analysis of this transcriptional dataset uncovered clear tends in gene expression characteristics for this species and new genes and pathways that associated with variation in pathogenic processes. Direct investigation confirmed functional predictions for a number of new regulators associated with growth and filamentation, demonstrating the utility of these approaches in linking genes to important phenotypes.

## INTRODUCTION

*Candida albicans* resides within the oral cavity, gastrointestinal tract, genitourinary tract, and on the skin of its human host as a commensal species (1). Development of an immunocompromised state can lead to *C. albicans* overgrowth of these same niches, producing debilitating mucosal infections and life-threatening bloodstream infections (2, 3). Critical to its success as both a ubiquitous commensal and opportunistic pathogen of multiple body sites is the ability for *C. albicans* to persist and proliferate in a wide range of physiological temperatures, oxic environments, nutrient availabilities, and pH conditions (4–6).

Clinical isolates of *C. albicans* represent a genetically diverse collection of heterozygous diploid organisms that can be separated into seventeen clades by multilocus sequence typing (MLST), with Clade I clade making up the majority of typed isolates (7–9). Recent sequencing efforts have examined genomes from across the *C. albicans* phylogeny (10, 11). Analysis of these genomes supports a primarily clonal lifestyle for *C. albicans,* with occasional inter-clade mating generating recombinant genomes in a subset of isolates (10, 12). Thus, *C. albicans* evolves principally through the acquisition and accumulation of iterative mutations, leading to expanded genotypic diversity over time.

This genotypic diversity contributes to extensive phenotypic variation among *C albicans* isolates, including an assortment of alternative cell states associated with distinct colonization and pathogenic traits (11, 13–21). Some phenotypes are biased towards specific *C. albicans* clades (22, 23). For example, inherent resistance to the antifungal 5-flucytosine (5-FC) is mediated by a single missense mutation in *FUR1* found ubiquitously across Clade I strains but absent in those from other clades (24, 25). In contrast, most phenotypes are heterogeneous both within and across *C. albicans* clades (11, 26–28), suggesting multi-locus control of these traits. This incongruence between genetic and phenotypic similarity in *C. albicans* deviates significantly from other asexual species in which phylogenetic conservation has been used to predict phenotypic traits (29–31). It has also complicated large-scale investigations of the underlying polymorphisms that contribute to *C. albicans* phenotypic diversity and limited identification of genotype-phenotype relationships (10, 11, 23). Instead, phenotypic diversity may associate more strongly with other molecular signatures such as gene expression and protein abundance (32–34).

The ability to rapidly respond to environmental cues is central to microbial adaptive potential. *C. albicans* adopts distinct transcriptional profiles in different cell states or when cultured in different physiologically-relevant conditions (13, 35, 36). Altered transcriptional states can be detected as early as 5 minutes following exposure to new environments (37–40). Distinct transcriptional responses in *C. albicans* are also observed in response to cues in the host, and these may contribute to colonization and pathogenesis in different niches (41–43).

Altered expression of hundreds of genes following environmental shifts complicates distinguishing the regulatory genes that govern these transcriptional changes from downstream effectors. Defining the genetic regulons associated with specific transcription factors or responses has typically relied on a simple model of conditional expression focused on a single gene or environmental condition (44–46), while the broader transcriptional architecture of *C. albicans* cells remains largely undefined. Concerted efforts to determine the transcriptional regulation of phenotypic switching between the *C. albicans* ‘white’ and ‘opaque’ states or between planktonic and biofilm communities has revealed the existence of highly interconnected transcription factor networks that collectively control differentiation between these states (47–51). Genes within these circuits encode some of the most well-characterized transcription factors in *C. albicans* and yet account for only a small fraction of the complete repertoire of transcriptional regulators. Thus, integration of large-scale expression data across *C. albicans* isolates could aid in elucidating the transcriptional networks underlying the regulatory architecture of this important human pathogen.

Here, we describe transcriptional profiling of 21 *C. albicans* isolates representing five clades with significant genotypic and phenotypic diversity (11). Gene expression profiles of these strains did not reflect their phylogenetic relationships at either the strain or clade level. Moreover, differential gene expression of up to 35% of the annotated genes was found between any two strains grown under identical conditions, with several strains displaying extensive strain-specific gene expression. Transcriptional differences between strains were associated with specific phenotypes that corroborate previous experimental studies and also predicted new molecular functions related to pathogenesis. Furthermore, unbiased clustering of genes based on correlated gene expression levels revealed a transcriptional map of cellular functions from which co-expression modules were linked to pathogen-associated phenotypes. Experimental investigation of two co-expression modules uncovered new regulators of filamentation and a cell state-specific module, and that these contribute to intra-species phenotypic variation in *C. albicans*.

## RESULTS

A previous investigation sequenced the genomes of 21 *C. albicans* isolates and identified widespread genetic and phenotypic variation among the strain set (11). Candidate gene approaches identified one strain with a homozygous nonsense mutation in the transcriptional regulator *EFG1* that caused a defect in filamentation and increased commensal fitness while decreasing systemic virulence (11). More recently, loss of *EFG1* function was also linked to formation of the “gray” phenotypic state in clinical isolates (13). However, broader attempts to link genetic polymorphisms to phenotypic differences present a significant challenge as multiple loci may regulate a single trait. Consequently, many of the causative polymorphisms contributing to phenotypic variation remain unknown. To gain greater insight into the underlying basis of phenotypic diversity in *C. albicans*, we transcriptionally profiled the set of 21 isolates with diverse geographical origins, sites of infection, and clade designations within the *C. albicans* phylogeny (Fig. S1).

### Gene expression does not reflect genetic relatedness

To compare gene expression across the 21 isolates, RNA was harvested from cells cultured in rich media (YPD, 30°C) in exponential phase. Transcript abundances were averaged between biological duplicates and binned across the 6,468 genes. The largest fraction of transcripts in the SC5314 reference genome were present at low but detectable expression levels (10-100 transcripts per million (TPM)), although the number of genes within each expression range fluctuated considerably among strains (Fig S2A, Table S1). For example, P37037 expressed 25.3% of its genes at less than 10 TPM whereas this proportion increased to ∼50% in GC75. Differential binning of gene expression even occurred among strains in the same clade (e.g., compare Clade II strains P57072 v. P76067), suggesting that large changes in genome-wide transcript abundance exist even between closely-related strains.

To determine if gene expression patterns were reflective of genetic relatedness, hierarchical clustering of genome-wide TPM values was performed. Similarity in gene expression profiles failed to reproduce the genetic phylogeny of these strains when averaged between replicates (Fig. 1A) or as individual samples (Fig. 2B). Variability in low abundance transcripts was not responsible for obscuring phylogenetic similarity as none of the 50 genes with the greatest dynamic range in expression recapitulated the phylogenetic tree (Fig. S2C). In fact, averaged expression of only 0.5% of all genes (31 of 6468) associated with phylogenetic similarity, and these genes were functionally enriched for transcriptional regulation by glucose (Fig. S2D).

**Figure 1.**
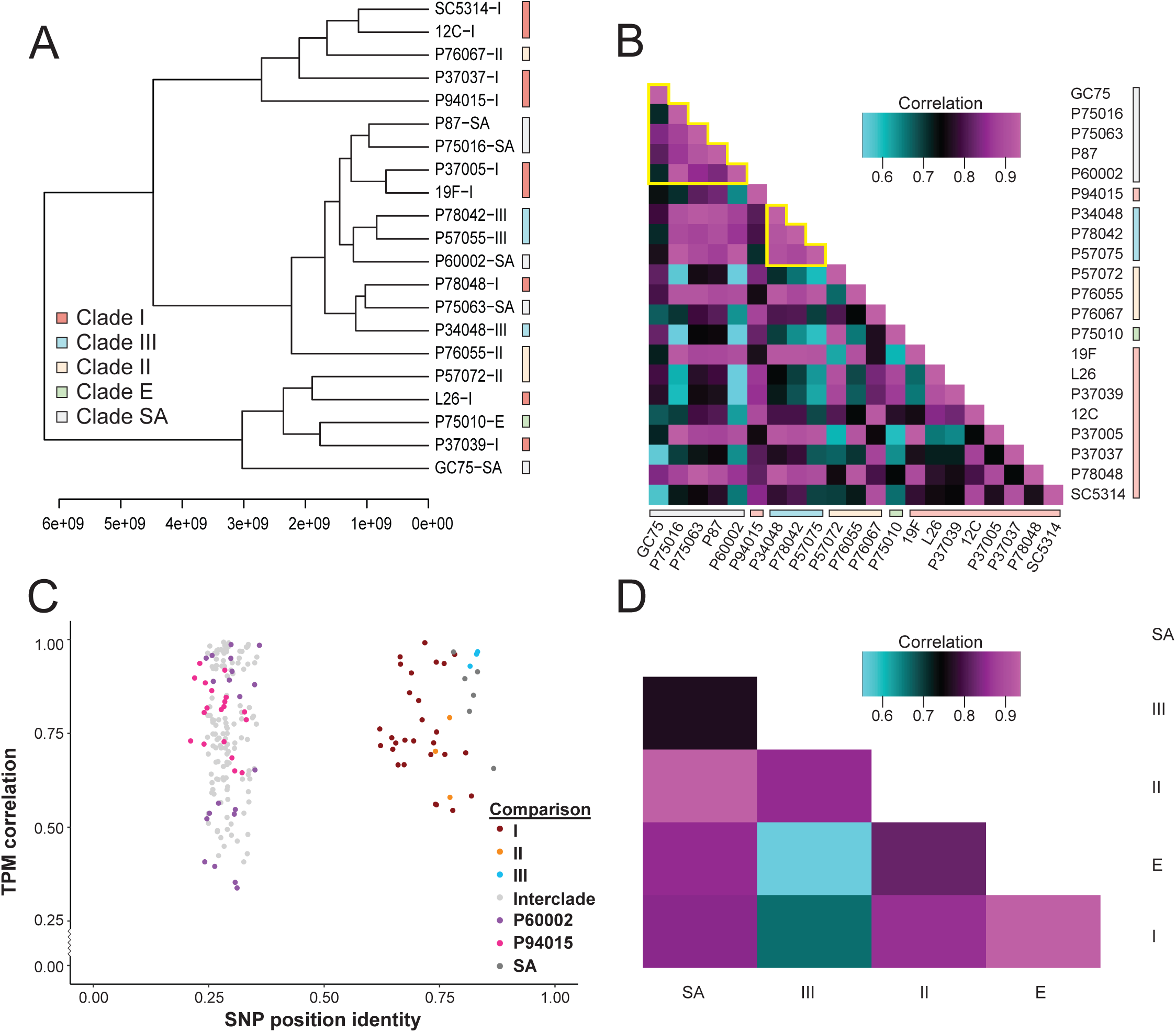
Gene expression does not reflect strain phylogeny. **A.** Hierarchical clustering of strains by Spearman’s correlation and average distances was performed for transcript abundance of the 6,468 genes across *C. albicans* strains using averaged values between replicates. Clade designation based on reported fingerprinting clades (FP) are indicated by color. **B.** Gene expression was averaged among biological replicates and the averages were compared between individual strains. Spearman’s correlation values were calculated in all pairwise combinations and visualized as a heat map ordered to reflect phylogenetic relatedness. FP clades are color coded and clades with strong clustering are outlined in yellow. **C.** The genetic similarity between isolates (x-axis) was compared to similarity in transcript abundance as defined in **B** (y-axis). Pairwise comparisons between all strains are represented as dots and color-coded to denote intra-clade comparisons (I-red, II-orange, III-blue, SA-dark gray) or marked as light gray for comparison across clades. Two clusters emerged with inter-clade comparisons showing less nucleotide similarity and a greater range of expression correlation scores (left) that extended below intra-clade comparisons (right). Two recombinant isolates, P60002 and P94015 (indicated in purple and magenta, respectively), clustered only within inter-clade comparisons. **D.** Clade gene expression profiles were built using the average of all strains within the clade. The clade-average profiles were compared by Spearman’s correlation and visualized by a heat map.

**Figure 2.**
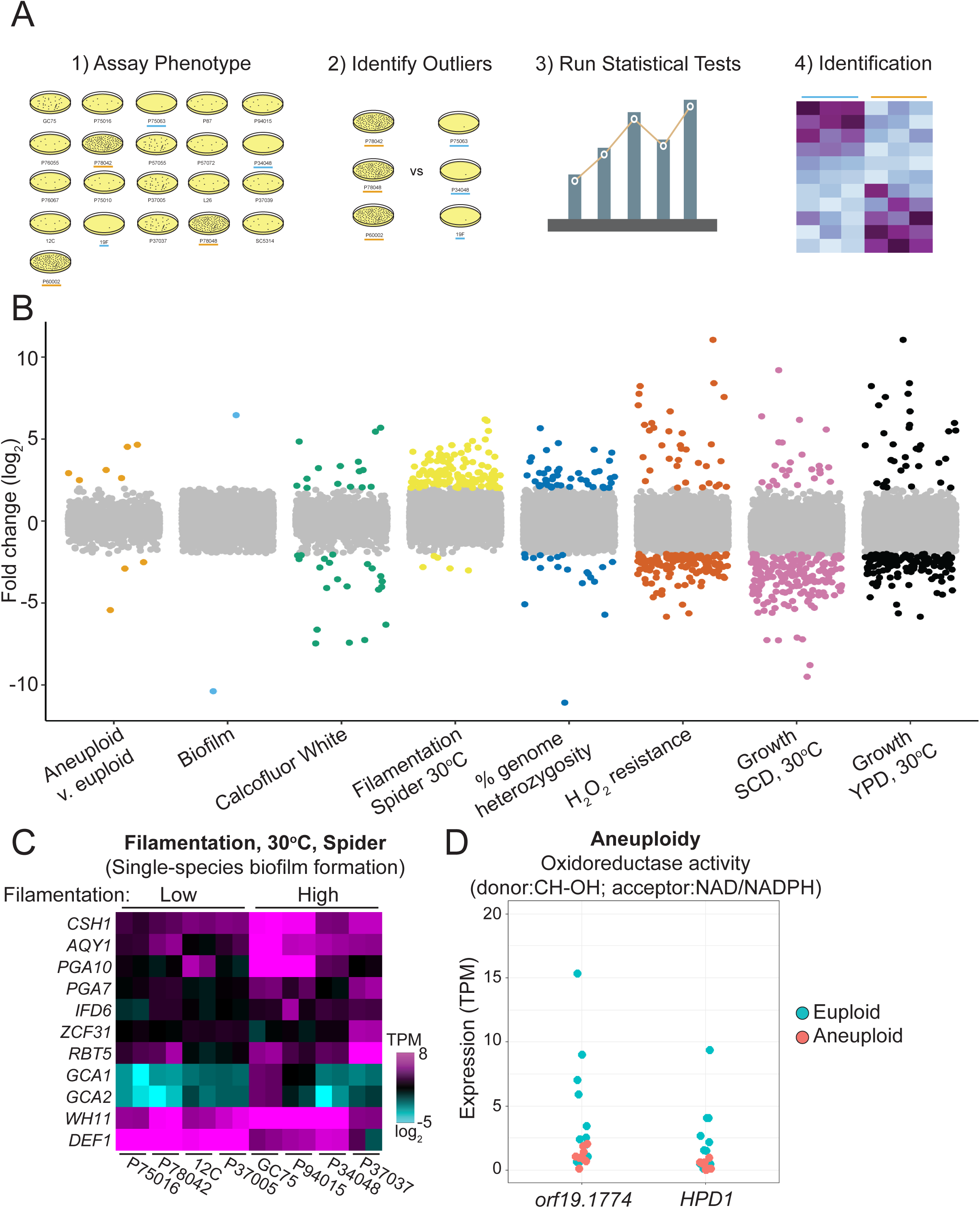
Differential expression predicts genes associated with *C. albicans* phenotypes. **A.** The workflow used to identify phenotype-associated genes is depicted. Phenotyping results for 8 traits determined in *Hirakawa et al.*, were used to: (1) screen strains to (2) identify strains with extreme phenotypes. (3) Differential gene expression (2-fold change, q<0.05) was identified among strains with opposing phenotypic groups, and (4) enrichment analysis performed for biological terms. **B.** The fold change in expression between groups with opposing phenotypic measurements as defined in **A** is plotted for all genes and the eight phenotypes investigated. Genes showing significantly different expression levels between the opposing phenotypic groups are color-coded by phenotype and genes without statistically supported differences are in gray. **C.** Transcripts per million (TPM) value are plotted as a heatmap on a log_2_ scale for differentially expressed genes within the enriched gene ontology term ‘Single species biofilm formation’ between strains that filament poorly (low) or profusely (high) on Spider agar medium at 30°C. Two biological replicates per strain are displayed. **D.** The TPM values for each euploid (blue) and aneuploid (red) isolate sample are plotted for the two differentially expressed genes within the enriched GO term for aneuploidy.

In a few select cases, averaged gene expression levels among strains within a single clade were similar, such as those within Clade III and among a subset of Clade SA strains (Fig. 1B, outlined in yellow). Indeed, gene expression within this strain set was more similar among intra-clade comparisons than inter-clade comparisons (Wilcoxon test, W = 4978, p-value = 0.022), supporting evidence of clade-associated expression signatures among these isolates. Regardless, intra- and inter-clade correlations of gene expression largely overlapped (average: 0.783 vs. 0.759; range: 0.33-0.97 v. 0.54-0.96, respectively, Fig. 1C), and Clade III strains largely drove the differences between intra-clade and inter-clade comparisons, which disappeared when these strains were removed (Brunner Munzel test (BM = 0.500, df = 78.9, p = 0.62). The two isolates in this set that have been proposed to harbor recombinant genomes, P60002 and P94015 (12), exhibited divergent genomes consistent with inter-clade comparisons of nucleotide divergence, and P60002 displayed the most divergent gene expression comparisons of any other strain (Fig. 1C). This further supports these isolates as being genetically distinct with unique expression patterns as compared to other strains from their assigned clades, and is in line with these two isolates having undergone inter-clade recombination during their evolutionary history (12).

Gene expression patterns were also compared between *C. albicans* clades, as some phenotypes have been associated with specific clades and clade-level comparisons can reduce the influence of ‘outlier’ strains (22, 23). However, with the exception of Clades II and III, similarities in clade-average expression levels were not enriched among the more closely-related clades (Fig. 1D, S3). Thus, genetic similarity contributes to, but does not strictly determine, similarity among *C. albicans* gene expression profiles.

### Gene expression differences between *C. albicans* isolates span biological traits

The set of *C. albicans* strains analyzed here exhibit up to 1.7% nucleotide divergence in pairwise comparisons (12), highlighting the potential for large-scale differences in genetic regulation and gene expression. The number of differentially expressed genes between any two isolates varied considerably, ranging from 43 to 1,457 genes (adjusted p-value 0.05, ≥ 2-fold change) (Tables S2, S3), and increased with greater dissimilarity in overall gene expression (Pearson’s test; r = -0.86, n = 210, p < 2.2E-16, Fig. S4). Investigation of gene ontologies (GO) associated with differentially expressed genes between isolates returned 147 process terms spanning the full breadth of biology (Table S4). The most prevalent GO terms were associated with ribosome biogenesis followed by nucleic acid and aromatic compound metabolism, suggesting that some isolates may have evolved unique growth characteristics, pathways to control nutrient utilization or signaling, and/or preferred nutrient conditions for optimal growth. Conversely, consistent gene expression levels across isolates point to core functions required for basic cellular processes in this diploid yeast. Of the 5,956 complete and intact open reading frames (ORFs) present across all 21 *C. albicans* isolates, 2,036 genes displayed indistinguishable expression levels among all strains. These include genes for key cellular functions such as amino acid charging of tRNAs, RNA polymerase function, and core translational processes (Table S5).

A distinctive class of genes considered were those expressed at unique levels within a single strain compared to all other strains and therefore classified as having strain-specific expression. The number of strain-specific genes varied considerably, ranging from 0 in isolates 12C, 19F, P37005, and P57055 to 171 in GC75 (q ≤ 0.05 and 2-fold change, Fig. S5). Strain-specific expression was enriched for cellular processes ranging from cell wall organization (GC75) to oxidation-reduction (P78042) to mannosytransferase activity (P60002) and RNA Polymerase I activity (P57072) (Table S6). Isolates with the largest number of uniquely expressed genes typically clustered closely with other strains in the phylogenetic tree (Fig. S1), further highlighting the disconnect between genetic relatedness and gene expression.

### Characteristics of functional noncoding RNA elements

Untranslated regions in *C. albicans* can serve as regulatory platforms for protein binding to control transcript stability and translation (52, 53). The average 5’ UTR length for 5,076 genes with detectable expression among all sequenced isolates centered at 1-25 bp and decreased in frequency with greater lengths (Fig. S6A). Prior analysis has revealed that some *C. albicans* transcription factors have extended 5’ UTRs greater than 1 kilobase (kb) in length (36, 52–54). Analysis of all transcription factor genes among the 21 strains showed they encoded significantly longer 5’ UTRs compared to the genome-wide average (286 v 97 bp, respectively; Wilcoxon test, W = 3.97E5, p-value < 2.2E-16; Fig. S6B, Table S7). In contrast, 3’ UTRs were, on average, between 25 and 75 bp for the 5,899 genes with detectable expression. Genes involved in protein translation were found to contain significantly longer 3’ UTRs than the genome average (141 v 44 bp, respectively; Wilcoxon test, W = 8.63E5, p-value < 2.2E-16; Fig. S6C, Table S8), which may also implicate important regulatory functions for these regions through either transcriptional or translational control (55).

Mobile genetic elements play an important role in shaping genome evolution through promoting recombination, disrupting gene function, and forming new transcriptional units (56). Previous work has catalogued the transposable elements (TEs) present in the *C. albicans* genome using their associated long terminal repeats for classification among clinical isolates (11, 57). Transcriptional profiling of the 21 *C. albicans* isolates revealed active expression of multiple transposon families within *C. albicans.* The most highly transcribed transposons were flanked by gamma class *LTR* sequences although the abundance of actively transcribed retro-elements varied immensely between strains (Fig. S7A). The RNA abundance of TEs did not reflect strain relatedness or changes in genomic copy number among the isolates (Pearson’s test; r = 0.062, df = 19, p = 0.79, Fig. S7B), suggesting that mechanisms of transposon quiescence or inactivation may contribute to differences in expression among strains.

### Gene expression does not correlate with chromosomal position

A previous report suggested that genes encoded at the chromosome ends could exhibit higher levels of expression plasticity, variable gene expression among cell populations (58). To assess expression plasticity, the coefficient of variation (CV) between biological replicates was calculated for all genes and averaged across the 21 strains. The average CV in 10 kb sliding windows remained fairly constant across the genome, centered at approximately 0.15 (Fig. <u>S8A</u>). Subtelomeric genes in the 15 kb most proximal to the telomeric repeats did not show increased variability compared to the rest of the genome; in fact, the CV decreased slightly in the subtelomeres. Additionally, only two of nine *TLO* genes with transcript abundance data across all strains showed elevated plasticity compared to the genome average (Students t-test; p<0.05, Fig. <u>S8B</u>). Instead, the majority of genes with significantly elevated expression plasticity were scattered throughout the genome (Table S9).

### Differentially expressed gene sets associate with *C. albicans* phenotypes

Previously, the 21 sequenced *C. albicans* isolates were characterized for a diverse set of *in vitro* and *in vivo* phenotypic traits (11). Differentially expressed genes between groups with extreme phenotypes can infer the causative networks or pathways that are responsible for the divergent traits (Fig. 2A).

To identify genes that associate with quantitative phenotypes, we compared differentially expressed genes between strains that displayed phenotypic extremes in Hirakawa et al. (11). Overall, gene expression profiles between groups for any given phenotype were overwhelmingly similar, with the extreme groups differentially expressing between 2 and 209 genes for each phenotype (> 2x change, q ≤ 0.05. Table S10). Growth phenotypes were associated with the largest number of differentially expressed genes (Fig. 2B), which may reflect the conditions used for RNA isolation (logarithmic phase growth in YPD medium at 30°C). Genes involved in cell cycle regulation, lipid metabolism, and carbohydrate metabolism were overrepresented among those differentially expressed between strains with fast/slow growth rates. Surprisingly, phenotypes not directly linked to the growth conditions in which RNA was prepared also showed differential expression of genes enriched for associated biological processes (Table S11). For example, strains with contrasting abilities to filament on Spider medium showed differential expression of genes associated with biofilm formation (11 of 129, q = 7.78E-3) and oxidoreductase activity (8 of 129, q = 9.61E-3), even though they were grown as planktonic cells in YPD medium at 30°C (Fig. 2C). Interestingly, strains harboring supernumerary chromosomes differentially expressed genes involved in oxidoreductase activity using NAD^+^/NADH acceptors compared to their euploid counterparts (2 of 9, q = 3.07E-2). Thus, gene expression differences could be connected to a variety of phenotypes, even though cells were isolated from a single experimental condition. This analysis was limited to phenotypes with clear opposing differences, however, and suggested that more dynamic models of expression-phenotype relationships could identify additional loci responsible for phenotypic variation.

### Linear models link gene expression with variation in simple traits

The differential gene expression analysis described above relied on categorical definitions (such as phenotypic extremes) and therefore failed to acknowledge that gene expression and quantitative traits often fall along a continuum. To incorporate non-discrete values, gene expression and phenotypic measurements were fit to a linear model. A generalized least squares model of regression was used to account for the potential influence of population structure on gene expression among the 21 strains. Expression values for the ∼6,400 genes were plotted for all 21 isolates against a panel of 23 phenotypic measurements spanning growth rates, drug resistance, stress resistance, filamentation, and virulence, and significant associations identified (Table S12). Notably, growth rates correlated strongly with expression of a significant portion of the genome (e.g., expression of 1,879 genes correlated with growth rates in YPD medium at 37°C, Fig. 3A). Genes connected to growth rates across a range of conditions were often overrepresented for functions related to the cell cycle or cell division (Table S13). For example, increased growth rates in minimal, Spider, and SCD media at 30°C displayed a linear relationship with increased expression of genes overrepresented in the mitotic cell cycle (q < 1.40E-4) and spindle assembly (q < 0.05). This analysis also identified core regulatory processes associated with growth rates including expression levels of Mediator, a major transcriptional regulatory complex (59). Expression of Mediator subunits were overrepresented for growth rates in YPD at 30°C, χ ((1, N=1320)=9.48, p=2.07E-3, Fig. 3B).

**Figure 3.**
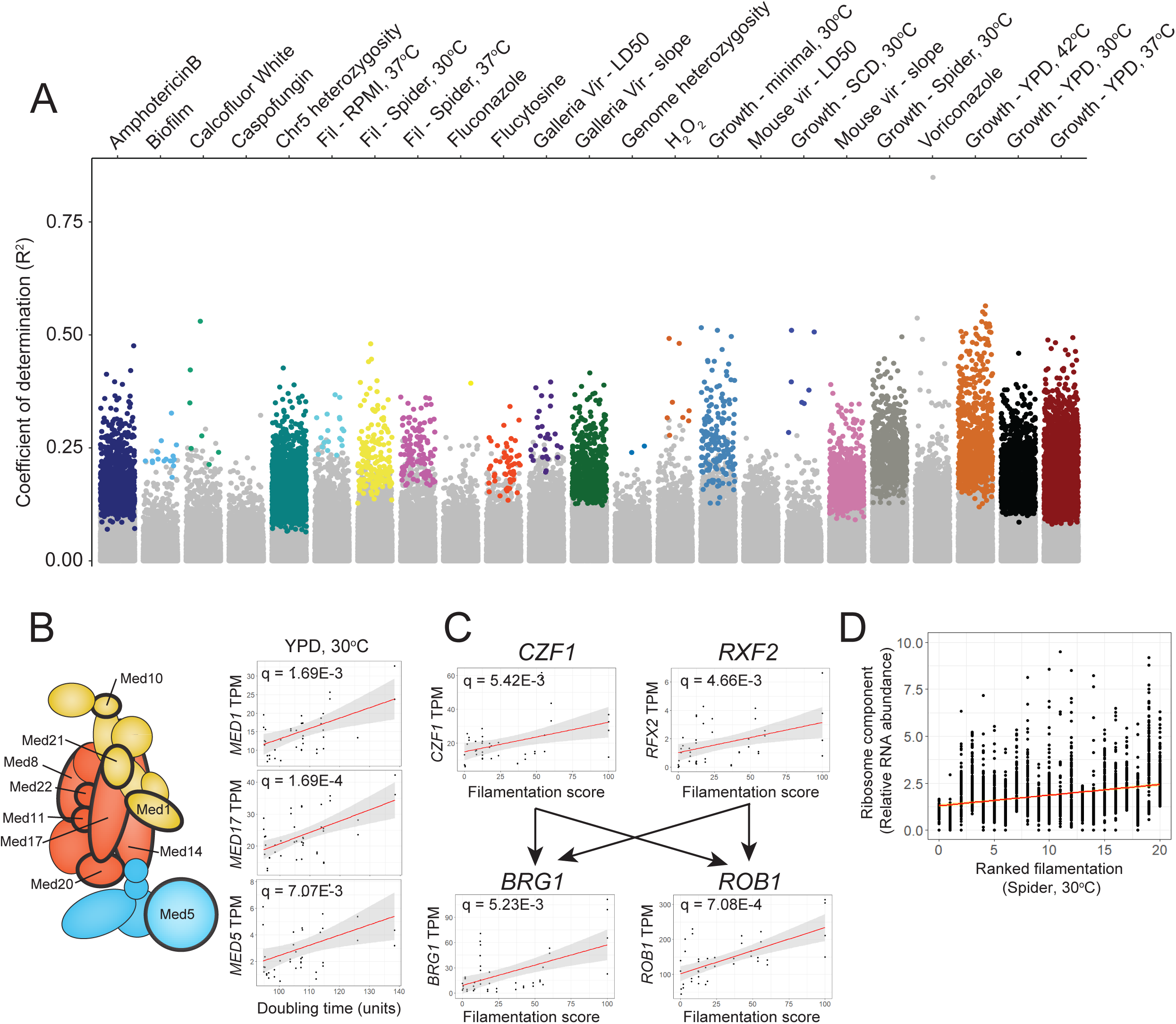
Linear regression reveals genes correlated with *C. albicans* phenotypic traits. **A.** Expression of each gene and quantitative phenotype scores from all biological replicates were fit to a linear model and tested for significance using Pearson’s correlation. The correlation score was plotted for each of 23 phenotypes and color-coded by phenotype for significantly associated genes. Gray points indicate no significant association. **B.** Representative correlation scores for components of the Mediator transcriptional regulator complex with growth in Spider medium at 30°C are indicated on the right. Mediator components significantly associated with these growth conditions are indicated in the Mediator schematic by thick black outlines. **C.** The expression of three genes previously known to be involved in *C. albicans* filamentation are plotted for the 21 isolates compared to their filamentation score on Spider solid medium at 30°C. The regulatory relationship of the three genes is indicated by arrows. *C.* The transcripts per million (TPM) value of all annotated ribosomal genes in the *C. albicans* genome is plotted for the 21 isolates by ascending filamentation scores on solid Spider medium at 30°C. A best fit line is indicated in red.

In contrast, linear modeling found fewer significant relationships between gene expression and more complex traits such as biofilm formation or virulence. Intriguingly, however, the expression of a large number of genes correlated linearly with the degree of hyphal growth observed in filamentation-inducing conditions. One of these genes, *CZF1*, is a key transcription factor required for the transition to hyphal growth (60), as well as a member of the core transcriptional network governing biofilm formation (47). Our results revealed that higher expression of *CZF1* in clinical isolates (in YPD medium) correlated with increased filamentation when cells were grown on Spider medium (Fig. 3C). Elevated expression of other hyphal-regulated genes including *RFX2, BRG1* and *ROB1* also correlated with increased filamentous growth under these conditions (q = 4.66E-3, 5.23E-3, and 7.08E-4, respectively). Both *BRG1* and *ROB1* are regulatory targets of Czf1 and Rfx2 (47, 51, 61), demonstrating that multiple members of known regulatory pathways can be uncovered by linear modeling of expression. Additionally, expression of ribosome and mitochondrial genes correlated with the extent of filamentation across a range of conditions (Fig. 3D), consistent with previous reports (62–64). Thus, linear modeling captured expression dependencies of key regulators with simple phenotypes but was less proficient in detecting relationships between gene expression and more complex *C. albicans* phenotypes.

### Construction of gene networks associated with phenotypic traits

To capture additional cellular pathways and processes associated with both simple and complex traits, we constructed gene expression networks using weighted gene correlation network analysis (WGCNA) (65). Implementation of network construction using transcript abundance of all genes across the set of 21 isolates produced 43 distinct co-expression modules (ME) (Fig. 4A, Table S14).

**Figure 4.**
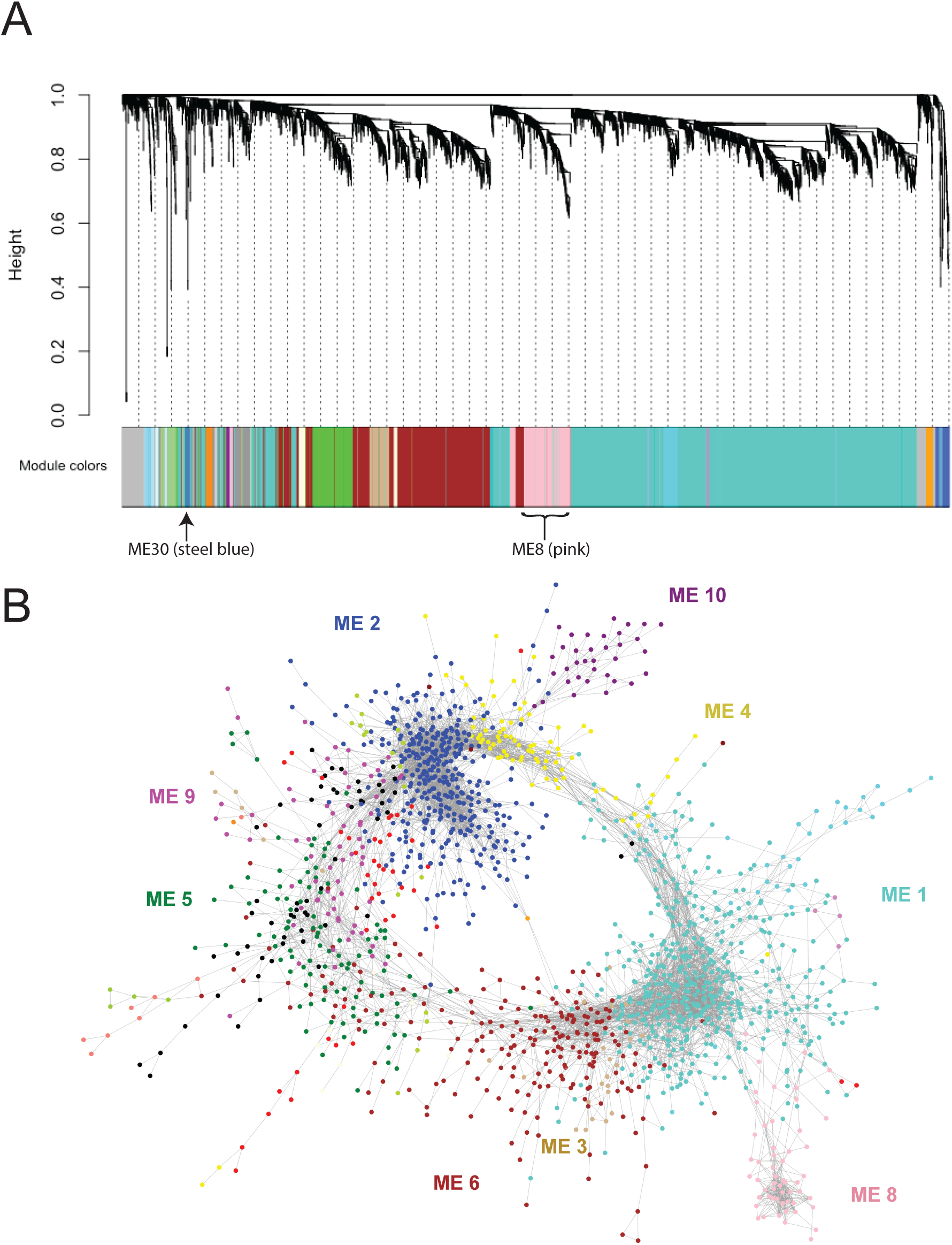
Co-expression modules reconstruct biological relationships in *C. albicans* cells. **A.** A weighted gene co-expression network analysis (WGCNA) of transcript abundance across all strains resolved 43 modules. A gene dendrogram obtained by average linkage hierarchical clustering is depicted above each associated module. ME8 and ME30 are indicated. **B.** The relationship between genes within all modules was visualized using a correlation cut-off of 0.93. Eight of the ten largest modules formed connections with each other and are color-coded as indicated. The relationship between each module is represented spatially where genes are represented as individual points and their correlated expression by edges.

Spatial organization of the co-expression modules produced a striking arrangement in which transcriptional crosstalk between modules was evident (Fig. 4B). Color coding was used to highlight different co-expression modules in which nodes are individual genes and edges have a correlation score of at least 0.93 (Fig. 4B). Surprisingly, we found that eight of the ten largest modules connect to one another to produce a ring structure, where most modules interact with a limited set of one to three other modules and that collectively incorporate expression of 67% of annotated *C. albicans* genes (4377 of 6468 genes). The two largest modules form the backbone of the ring structure: ME1 that includes the RNA processing and vesicular transport machinery, and ME2, which encompasses the translational machinery (Table S15). These processes are connected through ME4, which is enriched for genes involved in RNA binding in the nucleolus and ribosomal genes for RNA processing and translation. Genes required for ubiquitination and the proteasome are enriched in ME3 and connected to ME2, indicative of transcriptional crosstalk in protein turnover. ME3 is linked to ME5 that contains the genes for glycerophosphodiester transport and lipid production, to ME9, which is enriched for genes involved in the metabolism of nucleotide sugars and production of biofilm matrix, and finally to ME1, which links back to nucleotide processing in RNA metabolism. Thus, our analysis produced a gene expression atlas that delineates the interconnected transcriptional control of core cellular processes in *C. albicans*.

Gene co-expression modules were subsequently correlated to previously characterized phenotypes (11) to infer potential regulatory links (Fig. S9). Related phenotypes clustered to the same modules in many cases (e.g., growth rates in different media clustered to ME8, and filamentation across multiple conditions clustered to ME30). These module-phenotype links often included previously characterized genotype-phenotype associations. For example, elevated expression of ME30 and ME16 genes correlated with increased filamentation and encompassed known activators of filamentation such as *BRG1* (ME30) and *SUV3* (ME16) (66, 67). However, most genes in these modules have not been previously linked to filamentation and therefore represent candidates for further investigation.

### Identification of a putative state-specific network

Two phenotypes, growth rates and filamentation, were strongly associated with several gene co-expression modules (Fig. S9). To test WGCNA predictions of module-phenotype associations, we first interrogated the ME8 module, which was linked to growth rates under several conditions (Fig. 5A). Interestingly, a single strain, P37037, expressed genes in ME8 at higher levels than all other isolates (Fig. 5B), suggesting that ME8 conferred unique attribute(s) to this strain. The elevated expression of ME8 genes in P37037 may be due to coordinated gene regulation and/or interconnectivity, as 17 of the 18 genes within the ME8 network connect to a minimum of 12 other genes within the same network (Fig. 5C).

**Figure 5.**
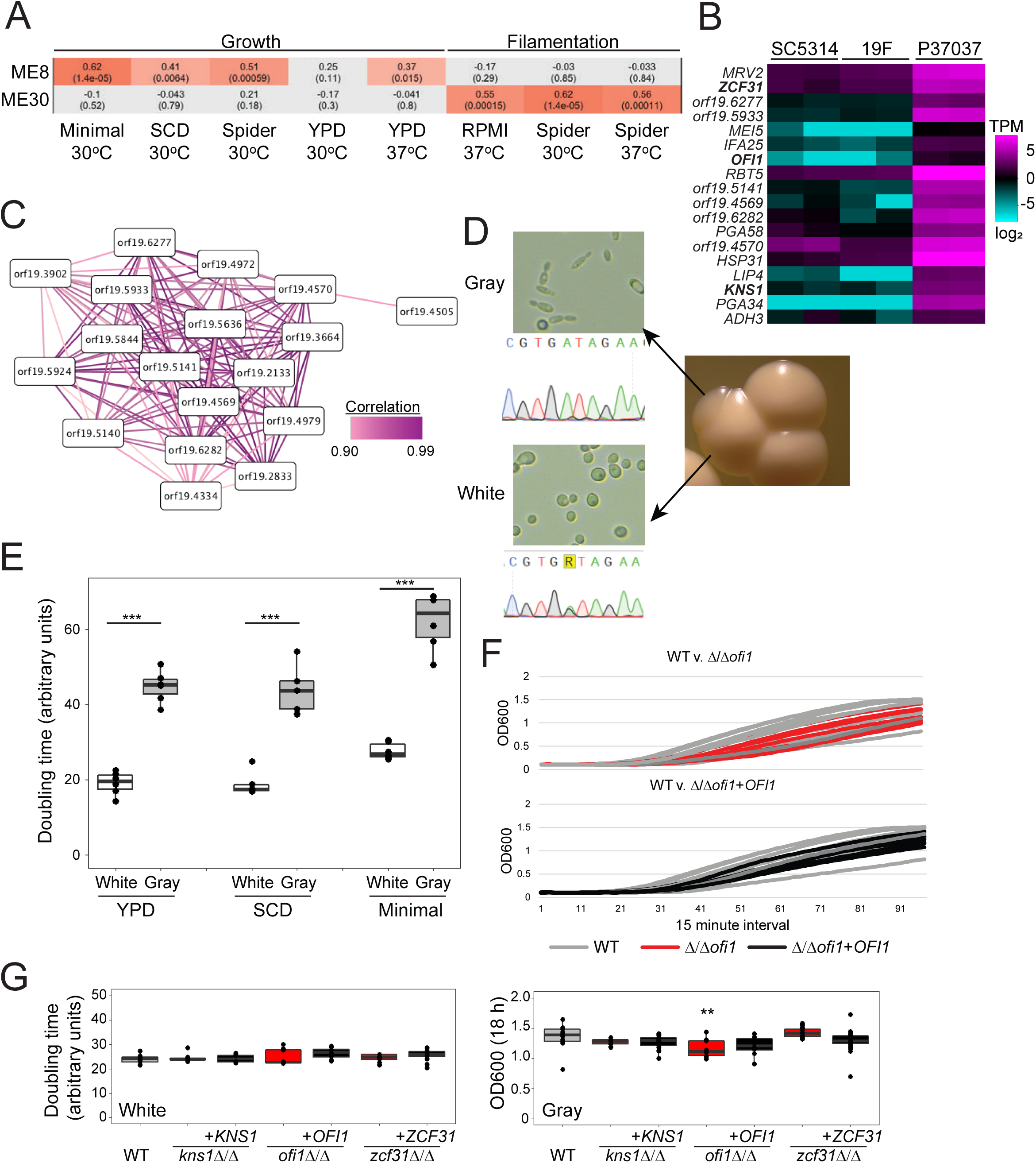
Identification of a gray-specific module associated with cell state growth differences. **A.** Two modules defined by WGCNA, ME8 and ME30, were correlated to phenotypes of the set of 21 *C. albicans* isolates. Significant associations are indicated by increasingly darker red hues and gray indicates no association. Each cell provides the Pearson’s correlation statistic (top) and q-value (bottom). **B.** A heatmap represents the transcripts per million (TPM) gene expression of ME8 genes on a log_2_ scale ranging from -6 to 6 for biological replicates for three isolates, SC5314, 19F and P37037. Genes in bold were tested experimentally. **C.** Strong correlated expression of 18 genes from ME8 is depicted where each gene is represented by modes and correlated expression shown as edges. Correlation scores are >90%. **D.** The white and gray cell states found in P37037 are shown for both colonies and cell images (at 40x magnification). The *EFG1* locus was genotyped by Sanger sequencing from both P37037 cell types. P37037 white cells encoded a heterozygous G/A and gray cells encoded a homozygous A/A at nucleotide 755 in *EFG1*. **E.** Growth rates for P37037 white and gray cell states. The average doubling time during logarithmic phase growth was determined in YPD, Spider, and minimal SD media and plotted as the mean with standard deviations. N=6. **F.** Growth curves during an 18-hour window are displayed for wildtype, Δ/Δ*ofi1*, and Δ/Δ*ofi1*+*OFI1* strains in the P37037 background and color coded as indicated. Measurements of optical density were taken in 15-minute intervals. **G.** Growth rates for white (left) and gray (right) cells in the wildtype, three mutant lines (Δ/Δ*kns1*, Δ/Δ*ofi1*, Δ/Δ*zcf31*), and their complemented P37037 strains. Significance was determined relative to the wildtype. N=6. ** denotes p<0.01. *** denotes p<0.001.

Analysis of P37037 colony sectors revealed two distinct cell types that resembled the previously defined ‘white’ and ‘gray’ states of *C. albicans* (Fig. 5D). *C. albicans* is most commonly isolated in the white state, which is considered the default state. In contrast, the gray state represents an *efg1/efg1* null state that can readily arise in strains that are *EFG1*/*efg1* heterozygous due to spontaneous loss of the functional allele (13). P37037 is functionally heterozygous for *EFG1* as it contains a polymorphism at nucleotide 755 that inactivates one allele via a G252D mutation in the encoded protein (13). Sequencing of the *EFG1* locus in P37037 confirmed the heterozygous polymorphic site (G/A) in white populations whereas all assayed gray colonies (4/4) had become homozygous (A/A) to produce cells lacking functional *EFG1* (Fig. 5D). Consistent with previous observations of conversion to the gray state (13), gray sectors often arose within white colonies but no white sectors were observed within gray colonies.

We hypothesized that gray cells within the mixed population from P37037 may be responsible for resolving the ME8 network and, potentially, its association with growth. Indeed, transcriptional profiling of gray P37037 cells demonstrated significantly elevated expression of ME8 genes compared to the white state (Fig. <u>S10A</u>). Interestingly, only 9 of these 18 genes displayed differences in expression between white and gray cells in the SC5314 background (Fig. <u>S10B</u>), indicating that strain background also influences white v. gray expression profiles. To test the association between cell state and growth, the doubling time of P37037 white and gray cells was compared in multiple media types. White cells grew significantly faster than gray cells in both nutrient-rich (YPD, SCD) and nutrient-poor (minimal) media at 30°C (Students t-test; p<0.001, Fig. 5E).

Three putative transcription factors in the ME8 module that had no previously described growth phenotypes (*KNS1*, *OFI1*, and *ZCF31*) were individually disrupted in strain P37037 to determine if genes within this module impact growth rates in either the white or gray cell state beyond the influence of cell state alone. Disruption of any of the three genes did not alter growth rates of white cells. In contrast, disruption of *OFI1* significantly decreased growth rates in the gray state, although doubling times were challenging to measure due to the lack of a clear logarithmic growth phase for these cells (Wilcoxon test; W(70), p = 0.017, Fig. 5F,G, <u>S11</u>). Loss of *KNS1* also decreased the growth rates of gray cells but this difference did not reach statistical significance (Fig. S11). Thus, genes in the ME8 module exhibit state-specific expression that reflects differences in growth between white/gray states.

### Dissection of a novel network that regulate filamentation

We also examined a second co-expression module, ME30, given that this module was associated with filamentation, but not growth rates, across a range of conditions (Fig. 5A). In contrast to ME8, this module displayed relatively low interconnectivity and exhibited a range of expression values across isolates (Fig. <u>S12A</u>). Expression of genes in the ME30 module was elevated in strains with higher filamentation scores compared to those that filament poorly (e.g., SC5314 v. P37037, respectively; Fig. 6A). ME30 genes included the previously characterized *BRG1* gene that encodes a transcriptional activator of filamentation (66), further suggesting a role for ME30 in promoting hyphal formation. Four genes from ME30 with potential regulatory roles (*UME7* – transcription factor, *FGR2* – putative transmembrane transport, *PHO100* – putative phosphatase, and *orf19.6864* – putative ubiquitin ligase), in addition to *BRG1*, were disrupted in the high expression strain SC5314 and assessed for filamentation in liquid and on solid media. Loss of each gene reduced filamentation in liquid RPMI medium at one hour, when hyphal initiation begins in SC5314 (Fig. 6B). Thus, most cells in the Δ/Δ*brg1* background remained as yeast whereas loss of the other four ME30 genes produced a heterogeneous mix of yeast cells and cells forming germ tubes. After four hours in RPMI media, all ME30 mutant cultures contained mostly hyphae although significantly fewer filamentous cells were present in the Δ/Δ*brg1,* Δ/Δ*fgr2*, Δ/Δ*pho100*, and Δ/Δ*ume7* strains (Wilcoxon test; p < 0.05, Fig 6B). Many of the mutants that formed filamentous cells remained as pseudohyphae at these later time points, compared to the wildtype background, which grew as a mix of hyphal and pseudohyphal cells (Fig. <u>S12B</u>). Complementation of each mutant restored the wildtype phenotype at both the one- and four-hour time points (Fig. 6B, <u>S12B</u>). Plating cells to single colonies on YPD and Spider media at 30°C produced similar outcomes with reduced filamentation of most ME30 mutants. Strains lacking *BRG1*, *FGR2*, and *UME7* demonstrated reduced colony filamentation after seven days on both YPD and Spider media with Δ/Δ*pho100* colonies also generating less filamentation on Spider medium (Wilcoxon test; p < 0.05, Fig. 6C). Similar to liquid filamentation, complementation of each mutant with a wildtype copy of the disrupted gene restored filamentation to wildtype levels (Fig. 6C). These results suggest that ME30 genes are responsible for activating filamentation responses in *C. albicans* and may be particularly important for hyphal initiation. Mutants in ME30 genes did not display any growth phenotypes, consistent with these defects being filamentation specific (Fig. 5A, <u>S12C</u>). Thus, our collective experimental validation of phenotypes predicted to associate with co-expression modules demonstrates the power of this approach to define gene function across *C. albicans* strains and to link previously uncharacterized loci to biological processes important for disease.

**Figure 6.**
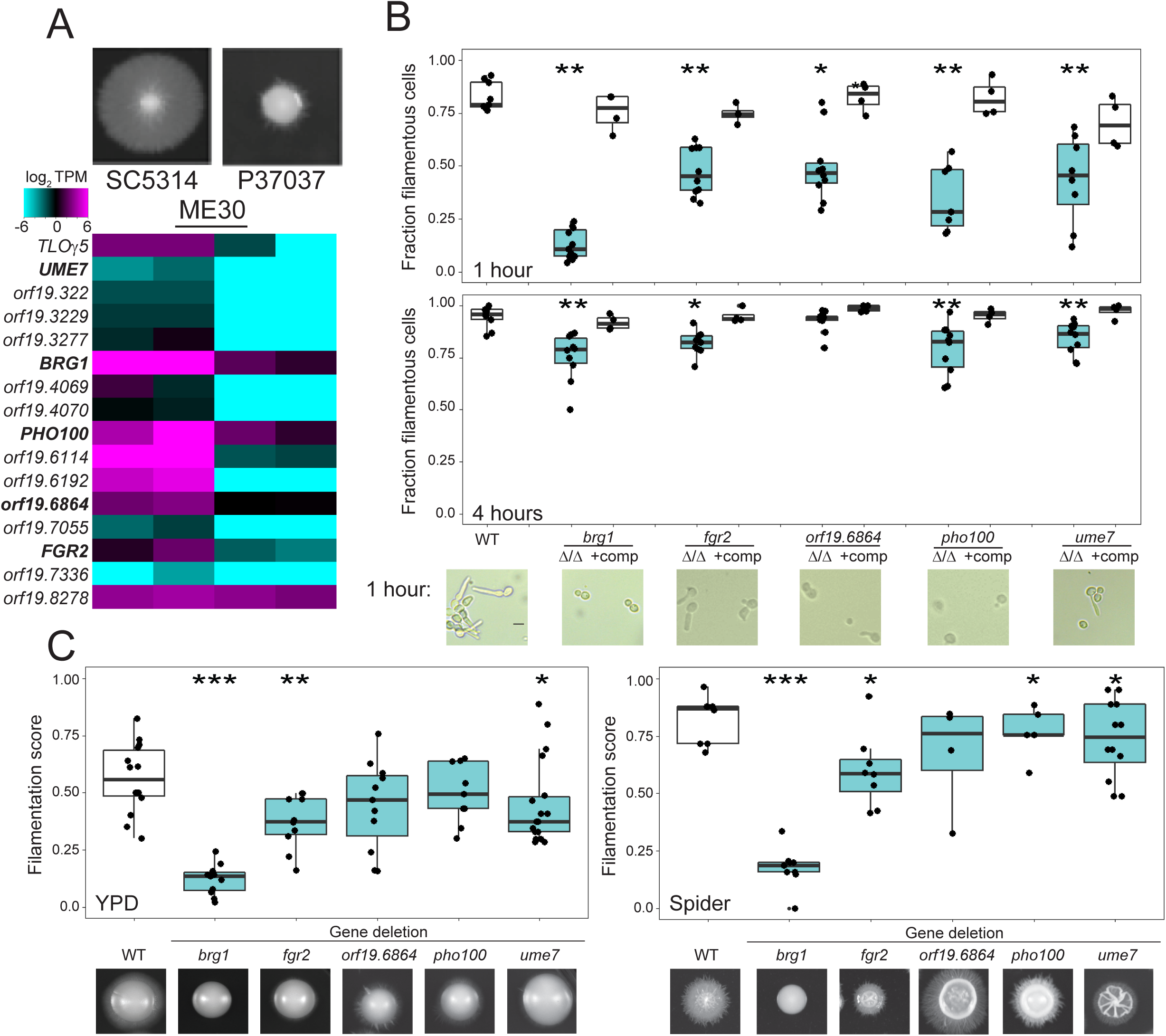
Genes within a co-expression module promote *C. albicans* filamentation across conditions. **A.** A heatmap represents the RNA transcripts per million (TPM) of all ME30 genes on a log_2_ scale ranging from -6 to 6 for SC5314 and P37037, isolates that filament strongly and poorly across multiple conditions, respectively. Genes in bold were tested experimentally. Colony images were taken following growth on Spider agar medium at 30°C for 7 days. **B.** SC5314 wildtype cells, mutants in five genes from the ME30 module, and the complemented mutants were grown for one and four hours in RPMI at 30°C and visualized at 40x magnification. Scale bar = 5 microns. **C.** The fraction of filamentous cells are plotted for SC5314 wildtype cells, mutants in five genes from the ME30 module, and the complemented mutants. N = 9,11, 4, 10, 4, 10, 4, 7, 4, 8, 4 for one hour and N = 9,10, 4, 10, 4, 10, 4, 10, 4, 10, 4 for four hours in order left to right. **D.** The filamentation score for SC5314 wildtype, ME 30 mutants, and the complemented mutants following growth on solid YPD (left) or Spider (right) media for seven days. N = 14, 14, 6, 14, 7, 13, 7, 9, 6, 19, 7 for YPD and 17, 12, 6, 12, 6, 8, 7, 8, 7, 14, 6 for Spider for strains from left to right. Significance was determined relative to the wildtype. * denotes p<0.05. ** denotes p<0.01. *** denotes p<0.001.

## DISCUSSION

A hallmark of *C. albicans* biology is the extensive genetic and phenotypic plasticity displayed among clinical isolates. This study expands previous observations that considerable transcriptional variation exists between natural isolates of the species (23, 27). We demonstrate that phylogenetic relationships between a set of 21 strains are not mirrored at the transcriptional level, as closely-related strains often display contrasting expression profiles under identical growth conditions. Notably, the construction of co-expression modules identified genes and pathways that underlie phenotypic differences between isolates. Furthermore, it permitted the direct evaluation of target genes for their roles in virulence-associated traits, thereby demonstrating the utility of this unbiased approach for delineating genes contributing to phenotypic diversity.

A striking finding in our analyses was the incongruence between constructed phylogenies and transcriptional profiles in *C. albicans*. Previous work has described transcriptional profiles in bacteria that reflect strain phylogeny and even phenotypic similarity based on shared lifestyle characteristics (68–70). In some eukaryotes such as *S. cerevisiae*, strong selective pressures based on niche specificity may explain incongruence between genetic and transcriptional profiles (34, 71). Here, we show that *C. albicans* strains express genes largely independent of their genetic similarity and that there is no clear association with the niche of isolation, although we recognize the limited number of multi-locus sequence type (MLST) clades represented by these isolates (7 of 17) as well as incomplete clinical information for these strains. The lack of a connection between genotype and gene expression is highlighted by the prevalence of strain-specific expression patterns for several isolates. This indicates that phenotypic variation between *C. albicans* isolates arises, in large part, from transcriptional differences that cannot be simply predicted by genetic phylogenies or clinical correlates.

Transcriptional differences among the 21 *C. albicans* isolates provided new insights into functional variation between isolates. Genes involved in metabolic processes were often differentially expressed among strains and may contribute to the range of growth rates seen for these isolates (11). Genes regulating transcriptional activation and hyphae formation also showed variable expression and were linked to differences in growth rates and filamentation, respectively. This is despite the fact that all expression profiling involved cells grown in a single culture condition (replete media at 30°C). Why might cells grown under one condition reflect expression differences that affect function in another? One possibility is that strains express genes in preparation for exposure to a new environment. Such priming can result from epigenetic reprogramming following a previous exposure (72), stochastic expression of regulators that promote bet hedging (73), and/or chromatin remodeling that favors activation of certain promoters (74). Priming of *C. albicans* cells could promote population fitness during environmental shifts including transitions between different host niches (75). *C. albicans* strains may also contain subpopulations of cells with distinct expression profiles that favor alternative environmental conditions, with the fraction of these subpopulations varying between strains. Additionally, cell variation in a population can arise due to changes in transcription factor binding that will disproportionately affect gene expression but will not cause general fitness defects (76). Single cell analysis and transcriptional profiling of large strain sets grown in multiple environmental conditions will help differentiate between these possibilities.

Our expression analysis of the set of 21 *C. albicans* strains facilitated the construction of a gene expression map of the species and the incorporation of a large proportion of uncharacterized loci into co-expression clusters linked to putative functions. Similar approaches in other systems have revealed the function of uncharacterized genes and their contributions to complex phenotypes (77–79). However, previous systems-level analyses have often skirted direct molecular testing of predicted gene functions. Here, experimental tests of *C. albicans* genes associated with growth and filamentation revealed functional roles for cell state and transcriptional regulators linked to two co-expression modules, ME8 and ME30. Analysis of genomic sequences could not predict the results described here as no inactivating mutations are present within ME8 and ME30 genetic alleles assayed in our strain set (11). Our study therefore reveals how expression profiling allows for an analysis of genotype-phenotype relationships using a variety of gene expression models instead of only assessing discrete mutation types.

Expression of ME8 module genes were linked to the gray cell state, which was recently shown to arise due to mutations that abolish *EFG1* function (13). The *EFG1* locus is heterozygous in P37037 and loss of heterozygosity (LOH) events can therefore cause cells to become *efg1* null and adopt the gray state (13). Unexpectedly, our analysis identified ME8 as a gray-specific co-expression module in P37037, where gray cells grow more slowly than white cells and which produced the expression module-phenotype association. ME8 genes that are upregulated in P37037 gray cells versus white cells are not uniformly upregulated in SC5314 gray cells (Fig. <u>S10B</u>). These results further emphasize that *C. albicans* phenotypes and expression profiles are dependent on their genetic background (11, 23, 26, 27). The existence of an *EFG1* heterozygote capable of accessing the gray state is not particularly uncommon (∼2% of assayed clinical isolates) and this hemizygous state may reflect advantages in gray state colonization of the gut or oral cavity compared to white cells (13, 15). Reduced growth rates of gray cells compared to white state cells in our assays could reflect differences from conditions in the host or, more simply, differences between genetic backgrounds. We evaluated the phenotypic consequences of deleting three genes from the highly interconnected ME8 module and showed that loss of *OFI1* significantly reduced the growth rates of P37037 gray cells. Thus, we uncovered a novel factor with a cell state-specific phenotype which further validated our approach.

A functional dissection of the ME30 module similarly connected several poorly characterized genes to a key phenotype in *C. albicans.* In this case, novel regulators of filamentation were discovered despite the wealth of research into filamentation pathways in this species (21, 80–82). Most studies have focused on genetic dissection of filamentation in SC5314 and have relied on candidate gene or transcriptional profiling approaches. We note that our identification of ME30 genes as regulators of filamentation did not rely on the presence of ORF-inactivating mutations but on differential expression across isolates that correlated with filamentation responses. Inclusion of the well-characterized filamentation regulator *BRG1* (66) emphasized the potential for other ME30 genes to regulate filamentation. Indeed, all assayed genes in ME30 appear to promote this process, albeit to different degrees, which likely reflects the lack of highly interconnected expression within this module (Fig. S12). All mutants of ME30 genes disrupted hyphal formation at early time points suggesting that these genes play a critical function in hyphal initiation and operate across multiple conditions, even though the ME30 module was defined using cells grown in the yeast form. The priming of filamentation via ME30 genes is supported by defined roles for Brg1 in recruiting Hda1, a histone deacetylase that remodels chromatin at the promoters of hyphae-specific genes, and occludes Nrg1, a negative regulator of filamentation (66, 83). Elevated expression of *BRG1* during rich medium growth could reduce the activation time needed to transcribe *UME6* and other genes that promote filamentation, while maintaining a phenotypically yeast state. The particularly long 5’ UTR of *BRG1* may indicate complex regulation of this gene, including undefined molecular pathways that include other ME30 genes, especially those with clear regulatory capacities (e.g., *FGR2*, *PHO100*, *UME7*) (54, 84). Thus, our study indicates that ME30 module genes may play broad roles in the regulation of filamentation in *C. albicans*.

## METHODS

### Media and reagents

Yeast extract peptone dextrose (YPD) and synthetic complete dextrose (SCD) media were prepared as previously described (85). Spider medium was prepared (1% nutrient broth, 1% mannitol, 0.2% K_2_HPO_4_) and equilibrated to a pH of 7.4. Minimal medium was prepared as 0.17% yeast nitrogen base, 0.5% ammonium sulfate. YPD containing 200 μg/mL nourseothricin (Werner Bioagents, Jena, Germany) was used to select for nourseothricin resistant (NAT^R^) strains.

### RNA-Seq library preparation

Two independent cultures for each of the 21 clinical isolates were grown at 30°C in YPD overnight. Cultures were diluted 1:100 into fresh YPD and allowed to grow to an OD of 1.0. RNA was harvested from cells using a Masterpure Yeast RNA Purification kit (Epicentre, Madison, WI) and treated with DNaseI (Fisher Scientific, Hampton, NH). RNA quality was measured on an Agilent 2100 Bioanalyzer and RNA with RIN scores ≥7.5 used for constructions of sequencing libraries.

Poly-A RNA was isolated and used to construct strand-specific libraries using the dUTP second strand marking method (86, 87) as previously described (88). The 42 sequencing libraries were pooled and sequenced on the Illumina HiSeq to generate 151 base paired-end reads. To measure gene expression, reads were aligned to the *C. albicans* SC5314 reference genome. RNA-Seq reads were then mapped to the transcripts with STAR (version 2.0.9) (89). Count tables were generated with HTSeq (version 0.9.0 (90), and differentially expressed genes were identified using EdgeR (version 3.28.1) (91). RNA-Seq data is available online and links are provided in Table S11.

### FASTQ Processing and alignments

Sequenced reads were returned in FASTQ format, and quality score confirmed using FastQC. All 42 samples exceeded the minimum allowed Phred quality score (28) across all bases. An average of 8.1 million reads were obtained per samples. Reads were aligned using the Spliced Transcripts Alignment to a Reference (STAR) with the alignIntronMin and alignIntronMax parameters set to 30 and 1000 (92). Greater than 90% of reads mapped to defined genes (range 96-98%). All other parameters were executed with default values. For each gene, the number of aligned reads was calculated using htseq-count (90). Gene features were defined as those exon regions annotated in the SC5314 Assembly 21 features file (http://shorturl.at/hpGW3), for a total of 6468 features. These read counts per feature were normalized into TPM values, which can be publicly accessed here: https://goo.gl/PqgGtH. The RNA-sequencing library contained a known defect with strand orientation, where orientation was incorrectly denoted as opposite of actual designation. All analyses (including features count) had taken this into account and corrected for it prior to analysis.

### Hierarchical clustering of gene expression

TPM values for all *C. albicans* genome features from the Assembly 21 genome feature file were used to build dendrograms of similar gene expression. Hierarchical clustering was performed using Spearman’s correlation and average linkage. To assess, similarity between biological duplicates trees were built and tested with 1000 bootstraps using the ‘pvclust’ package (version 2.2-0) in R (version 3.5.3). For comparisons across strains, average TPM values were calculated between strains and hierarchical clustering performed.

### Correlation of expression with strain phylogeny

Phylogenetic relatedness among the 21 clinical isolates focused on strains that clustered well within their respective canonical clusters (I, II, III, SA). To increase the tightness of these well-represented clusters, outlier strains with long-branch lengths (P94015, P60002, and P75010) were removed. Based on each gene’s individual transcriptomic profile, we performed unsupervised clustering on each gene’s expression for the remaining 18 strains to bin into 4 groups using the R library kmeans. Hierarchical clustering was then performed on those genes for which these 4 groups contained at least half of the expected strains organized the same as for whole genome analysis. For each gene’s hierarchical clustering, the number of strains inconsistently assigned were counted and only 31 genes had at most six incorrectly assigned strains, less than expected by chance. No gene reported perfect homology with the phylogenetic tree.

### 5’ UTR and 3’ UTR construction

The aligned reads in bam file formats for each of the 42 replicates was converted into bed format using [bamToBed](http://bedtools.readthedocs.io/en/latest/content/tools/bamtobed.html), such that each individually aligned read is denoted in each row. Next, [mergeBed](http://bedtools.readthedocs.io/en/latest/content/tools/merge.html) was applied so that overlapping reads on the same strand are merged together into one contiguous segment. [IntersectBed](http://bedtools.readthedocs.io/en/latest/content/tools/intersect.html) was used to annotate the respective gene contained with each overlapping segment, with a minimum overlap of 1 bp. The -S flag was used when running [intersectBed](http://bedtools.readthedocs.io/en/latest/content/tools/intersect.html) to account for opposite strand orientation. Continuous merged reads that overlap with more than one gene feature and those with negative UTR lengths were removed.

### Differential gene expression by phenotypic extremes

Previous phenotyping of these 21 was used as the basis for this analysis (11). For each phenotype with categorical extremes, both biological replicates for strains exhibiting traits at the extremes of the distribution for each phenotype were binned into opposing groups and compared against each other for differentially expressed genes as described above using EdgeR (91). The following groupings were used for each phenotypic comparison:

a. SCD30°C: P60002, P78048, P37037 vs GC75, P75063, P34048 (slow vs fast)
b. YPD30°C: P76067, P94015 vs P34048, SC5314, P75016, GC75, P57055 (slow vs fast)
c. Biofilms: GC75, P87, SC5314 vs P75016, P94015, P57072, P75010 (heavy vs light)
d. FilamentationScoreSpider30: P75016, P78042, 12C, P37005 vs GC75, P94015, P34048, P37037 (high vs low)
e. CalcofluorWhite: GC75, P75016, P75063, P60002, P75010, 19F, L26, P37039, 12C, P78048, SC5314 vs P34048, P57055, P57072, P76055, P76067 (colonies at 4^th^ dilution versus the 1^st^ dilution)
f. HydrogenPeroxide: P75016, P75063, P87, P60002 vs P94015, P78042, P57055 (colonies at 4^th^ dilution versus none at any dilution)
g. GenomeHeterozygosity: P75016, P34048, P78042, P78048, SC5314 vs P87, P94015 (high vs low)

Differentially expressed genes were filtered for a minimum log_2_ fold change of 2, a qValue less than or equal to 0.05, and included only genes that had a minimum of 1 count per million reads in at least two samples. The expression dataset was normalized using the default weighted trimmed mean of M-values (TMM) method and dispersion estimated using an empirical Bayes method. Because all replicates were collected and sequenced in a single experimental run, no batch effect is expected.

### Gene ontology annotation

Enrichment for gene ontology terms was conducted through the Candida Genome Database (93). In complement, we introduce an R library (CAlbicansR) to facilitate non-browser analysis of *Candida* genomic datasets. Its functionality includes an offline database for converting orf19 identifiers into gene names, and vice versa. In addition, the library also provides a function for automated searches of the Gene Ontology Term Finder. Results are outputted into the R console.

### Linear regression of phenotype on gene expression

The strength of a linear association between a gene’s expression and phenotypic score was assessed for all genes in all phenotypes using each sequencing set as a single data point (42 data points in all). To account for existing phylogenetic relationships, the covariance structure between strains was calculated based a Brownian motion process of evolution, using the R phytools package. Phylogenetic generalized least square regression was fitted while accounting for within-group correlation structure as defined previously. For each gene, the x-axis represented the strain’s expression of that gene and the y-axis indicated the corresponding strain’s phenotype score, and a linear least-squares equation was calculated. The F-statistic was used to assess statistical significance, with a Bonferroni correction applied to each set of phenotype tests. Only genes with a corrected p-value less than 0.05 were retained.

### WGCNA construction

The recommended default settings were used from the [tutorial](http://labs.genetics.ulca.edu/horvath/CoexpressionNetwork/Rpackages/WGCNA/Tutorials/FemaleLiver-02-networkConstr-auto.pdf) section 2.a.2 for WGCNA of all 42 sequenced samples (2 replicates each from 21 isolates). Specifically, beta was set to 20 to achieve scale-free topology (first value for which R_2_ exceeded 0.80) as recommended (94). In addition, the networkType and TOMType both were set to signed, minModuleSize at 10, and mergeCutHeight at 0.15.

### Identification of bimodal networks

To identify genes with expression values that follow a multimodal distribution, we used a [Gap Statistic](http://web.standford.edu/~hastie/Papers/gap.pdf) method implemented through the R library [clusGap](http://stat.ethz.ch/R-manual/R-devel/library/cluster/html/clusGap.html), and used [hclust](http://stat.ethz.ch/R-manual/R-devel/library/stats/html/hclust.html) to identify clusters. Only genes with minimum expression values were considered (TPM >= 5). A gene was considered to operate via a bimodal response if its maximized gap statistic exceeded 0.9 and corresponding k value exceeded a minimum of 2. Specifically, this analysis identified a subset of genes within ME8 that express significantly higher only in P37037.

### Strain and plasmid construction

Strains, oligonucleotides, and plasmids described in this paper are provided in Tables S16, <u>S17</u>, and <u>S18</u>, respectively. Gene disruption was performed using long oligonucleotide-mediated targeting of *OFI1*, *ZCF31*, and *KNS1* in P37037 through amplification of the *SAT1*-*FLP* cassette from pSFS2A (deletion oligonucleotides listed in pairs as “Round 1 KO” or “Round 2 KO” in Table S17) and integrated by lithium acetate transformation (95, 96). Integration of deletion cassettes (Deletion Chk) and complementation plasmids (Addback Chk), as well as the presence or absence of open reading frames for each gene (ORF Chk), were confirmed with PCR using the oligonucleotides listed in Table S17. The *SAT1-FLP* cassette was recycled by plating to 100 colonies on yeast extract peptone maltose (YPM) solid media top-spread with either 10 μg/mL or 20 μg/mL NAT. Small colonies were then patched to YPD with or without 200 μg/mL NAT to screen for nourseothricin sensitive (NAT^S^) colonies.

Construction of the *OFI1* complementation plasmid p41 was performed by cloning PCR amplified *OFI1* from P37037 genomic DNA (including the promoter, coding sequence, and downstream) into pSFS2A using restriction enzymes ApaI and BamHI. The resulting plasmid was linearized in the promoter of *OFI1* using HpaI for transformation into *C. albicans*. Construction of plasmids p50, p52, and p53 were performed using gap-repair cloning as described in *Jacobus et al.,* (97) to generate *ZCF31_A*, *ZCF31_B*, and *KNS1* complementation plasmids, respectively. Briefly, *ZCF31* from P37037 genomic DNA (including the promoter, coding sequence, and downstream) was PCR amplified with oligonucleotides encoding 20 bp ends homologous to pSFS2A, and pSFS2a was linearized via PCR amplification with oligonucleotides containing 20 bp of homology to *ZCF31*, generating 40 bp of overlap. After digestion of residual plasmid template using DpnI, each PCR product was gel purified and co-transformed into chemically-competent DH5α to be assembled into an intact plasmid. The resulting plasmids yielded two plasmids containing different *ZCF31* alleles listed as p50 (*ZCF31-*_P37037_A_) and p52 (*ZCF31-*_P37037_B_). p50 and p52 were linearized in the promoter of *ZCF31* using PacI for lithium acetate transformation into *C. albicans*. The *KNS1* complementation plasmid p53 was generated in a similar manner, but the genomic amplification was split into two fragments to introduce a novel MluI restriction site into the promoter region. p53 was linearized in the promoter of *KNS1* using MluI for *C. albicans* transformation.

Pure populations of P37037 white and gray state cells were isolated from the mixed P37037 stock by streaking MAY3 onto YPD and growing at 30°C for 5 days until individual white and gray colonies could be differentiated. Independent colonies were inoculated into liquid YPD and grown overnight at 30°C for storage and sequencing of *EFG1* to determine the allelic makeup of this locus.

Gray state cells from P37037-derived mutant strains were obtained by streaking white state strains onto YPD, followed by growth at room temperature. After five days of growth, gray sectors were identified and struck out onto YPD and grown at room temperature once again to obtain isolated gray state colonies. After three days of growth, streaks were examined at a cellular and colony level to confirm gray state morphologies.

CRISPR-mediated deletion of SC5314 *BRG1*, *UME7*, *orf19.6864*, *PHO100*, and *FGR2* were performed as previously described using a modified lithium acetate transformation protocol (98). Colonies were screened for gene deletions by PCR for the presence of a band using oligonucleotides flanking the excised locus (Up/Dwn Check) and for the loss of the target gene (ORF Chk) using the oligonucleotides listed in Table S17.

Complementation plasmids for *BRG1*, *UME7*, *orf19.6864*, *PHO100*, and *FGR2* mutants were constructed by amplifying the wildtype locus from the background strains for all CRISPR-based deletions using primers listed in Table S17 and cloning them into pSFS2a as described above using gap repair cloning. All genes were cloned in two pieces with the exception of *UME7*, which required a three-piece cloning to include a MluI site for linearization prior to transformation (plasmids listed in Table S18). Genes were confirmed to be identical to the expected sequence by Sanger sequencing and then linearized using PacI, MluI, PacI, AgeI, and CspCI for *BRG1*, *UME7*, *orf19.6864*, *PHO100*, and *FGR2*, respectively, for lithium acetate transformation. Cells were selected on 200 μg/mL NAT and confirmed to contain the gene integrated at the native locus by PCR using primers listed in Table S17.

### Filamentation

For liquid filamentation assays, cells were grown overnight in YPD at 30°C. Cultures the next day were spun down, washed in PBS, and inoculated 1:100 into RPMI 1640 liquid medium and allowed to grow for either 1 or 4 hours before imaging. Images were captured at 40x magnification across 6 fields of view per sample to include at least 50 cells. At least four biological replicates were performed per genotype.

For solid media filamentation, cells were taken from YPD solid medium, counted by hemocytometer, and plated to Spider or YPD at 100 cells per plate. Plates were incubated at 30°C for seven days and imaged. Filamentation was measured using MIPAR as previously described (99). At least six biological replicates were performed per genotype.

### Data availability

The datasets generated during and/or analysed during the current study are available from the corresponding author on reasonable request. The transcriptional profiling data generated in this study have been submitted to the NCBI BioProject database (https://www.ncbi.nlm.nih.gov/bioproject/) under accession number PRJNA630085. Tools developed to aid in gene ontology analysis are available from https://github.com/joshuamwang/CAlbicansR.

## FUNDING

This work was supported by National Institutes of Health grants R01AI148788 to M.Z.A. and R01AI141893/R01AI081704 to R.J.B. R.J.F. was supported by an NIH F31 fellowship (1F31DE029409-01). This work was also supported by the American Heart Association grant AHA 20PRE35200201, M.J.D. 2020.

## ACKNOWLEDGEMENTS

We thank the entire Anderson lab for helpful discussions and feedback during the production of this work. We also than the lab of Dr. Chad Rappleye for comments and critique of this work and Drs. Kou-San Ju, Christina Cuomo, and Lara Sucheston-Campbell for feedback on network analysis and visualization.

---

**Supplemental Figure 1. Phylogenetic relationship of *C. albicans* strains used in this study.** The phylogenetic relationship of the 21 *C. albicans* isolates used for transcriptional profiling is shown based on comparison of full genome sequences. Bootstrap support for each node is indicated. Assignment of isolates to fingerprinting clades are color coded.

**Supplemental Figure 2. Correlation of gene expression with phylogenetic relationships among the *C. albicans* isolates. A.** Read counts were calculated for all genes from each strain and binned based on the transcripts per million (TPM) value. The fraction of reads within each bin was then plotted per strain. Clade assignments for each strain are color-coded as indicated. **B.** Similarity in transcript profiles among the 42 biological samples was assessed by hierarchical clustering of TPM values using Euclidean distance and average linkage. 1000 bootstraps were performed. The resulting bootstrap value are shown in green and corresponding approximately unbiased (AU) p-values are shown in red at each node. **C.** A heatmap represents the RNA transcripts per million (TPM) of the 50 genes with the greatest difference in expression among the 21 isolates on a log_2_ scale. The expression for each strain is the average of two biological replicates. The strains are ordered based on their phylogenetic relationships and their clade assignments are color coded. **D.** The 32 genes whose expression significantly correlated with the strain phylogeny are listed. Genes that contributed to enrichment of the gene ontology (GO) terms associated with this list are bolded. Significant GO categories are listed.

**Supplemental Figure 3. Transcriptional profiles are not more similar among genetically similar strains.** A distance matrix based on similarity in transcriptional profiles was constructed for all 21 *C. albicans* isolates. Distances were separated based on comparison between strains within the same clade or between strains in different clades based on fingerprinting analysis and plotted. Intra-clade and inter-clade comparisons were not statistically different.

**Supplemental Figure 4. Greater dissimilarity in gene expression correlates with more differential gene expression.** The number of differentially expressed genes between any two strains (adjusted p-value < 0.05, 2-fold cut-off) and the similarity in overall gene expression between two strains in all pairwise comparisons was plotted. Comparisons were performed in all pairwise combinations for all strains and color-coded for comparisons between two strains within the same clade or marked as gray for comparison across clades. This data produced an inverse relationship between expression similarity and the number of differentially expressed genes.

**Supplemental Figure 5. Strain-specific gene expression among *c. albicans* isolates. A.** The number of genes expressed uniquely by one strain compared to all other 20 transcriptionally profiled isolates were plotted for each of the 21 isolates. Isolates that uniquely expressed a greater number of genes beyond two standard deviations are labeled. **B.** The number of strain-specific genes for each isolate is listed.

**Supplemental Figure 6. Untranslated regions (UTRs) in *C. albicans* vary in length with gene function. A.** The UTR length for all genes in each isolate was determined by measuring the length of continuous reads extending beyond defined coding sequences on the appropriate strand. Lengths for each gene were plotted with 5’ UTRs above and 3’ UTRs below the x-axis. Red vertical lines indicate the 95% cutoff value. **B.** The 5’ UTR was detected from aligned transcripts from each of the 21 sequenced isolates. The length of the 5’ UTR for each gene was averaged for all genes with detectable expression in at least 15 strains. The length of all gene 5’ UTRs is plotted alongside those of all *C. albicans* transcription factors as defined in the Candida Genome Database (http://candidagenome.org). **C.** The 3’ UTRs of all genes in the *C. albicans* genome was similarly determined from transcriptional profiling. The 3’ UTRs of all genes was plotted alongside all genes defined by the gene ontology term ‘ribosome’.

**Supplemental Figure 7. Retroelement expression does not correlate with copy number. A.** The abundance of each transposon-associated long-terminal repeat (LTR) was determined from RNA-Seq for each strain and is shown as a stacked bar and color-coded to indicate each LTR class. Strains are color-coded by clade. **B.** The number of retroelements encoded in the genome of each *C. albicans* isolate was determined from previous whole genome sequencing (11), and plotted against the total transcripts per million (TPM) value for all retroelements. A linear model was fit to the data to detect a relationship between copy number and expression.

